## Supplementary for "Out of the single-neuron straitjacket: neurons within assemblies change selectivity and their reconfiguration underlies dynamic coding"

### Supplementary information

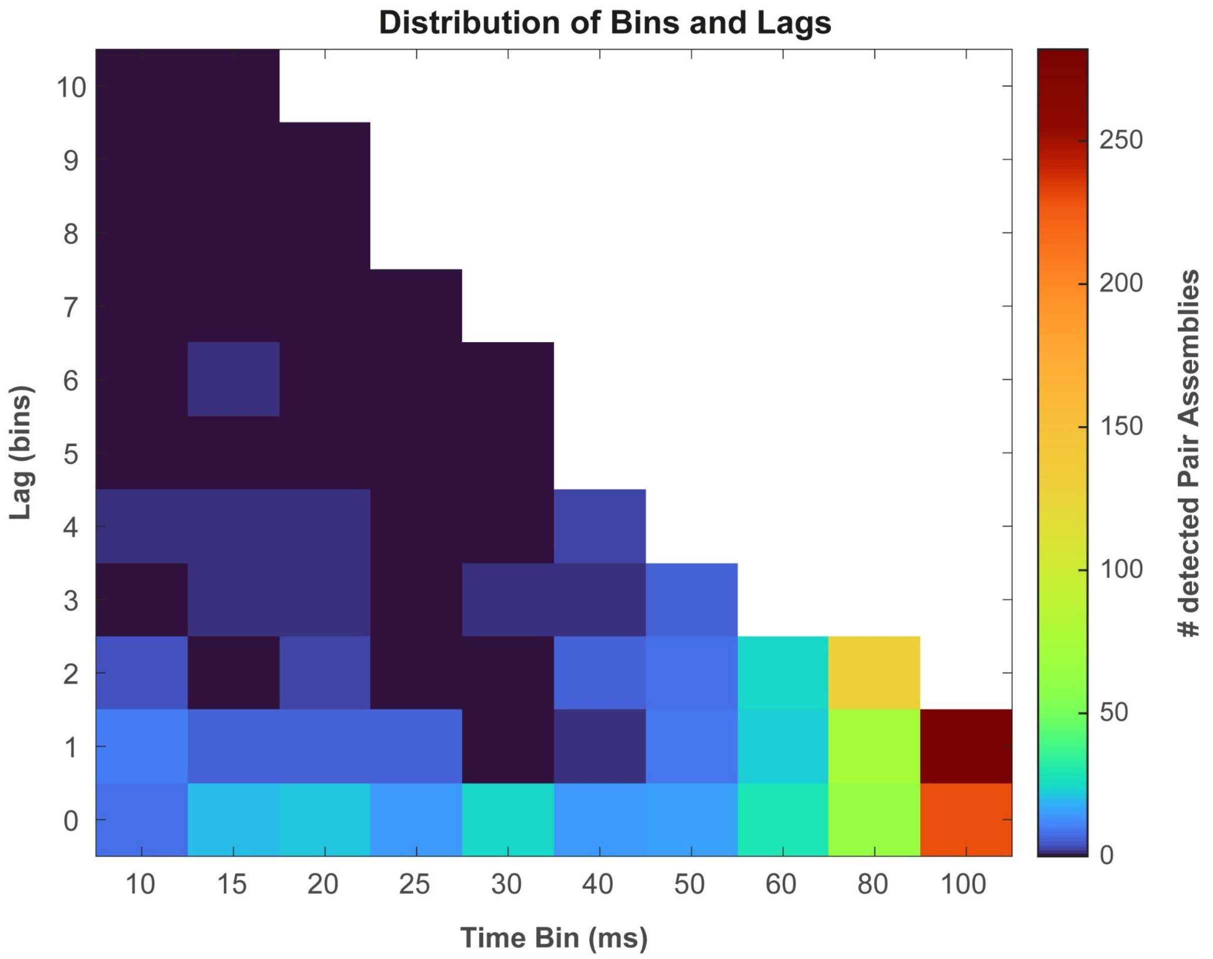


**Figure S1. Distribution of bins:** Heatmap of the distribution of bin/lag combinations for the pair assemblies identified by the algorithm in either task. *Bin* denotes the characteristic time scale at which the assembly was detected, while *lag* refers to the time latency, expressed in time bins, between the activation of the first and the second neuron in the pair assembly.

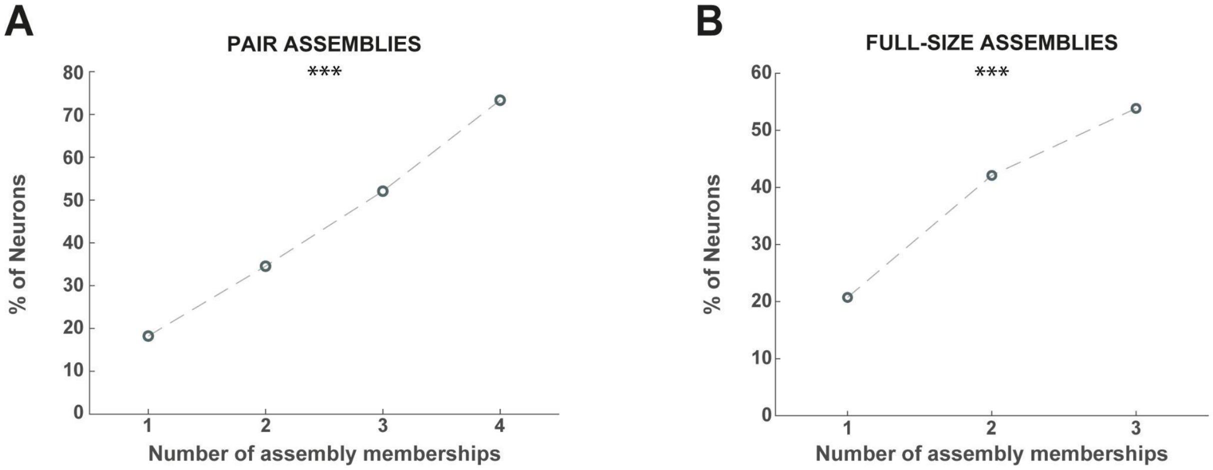


**Figure S2. Contribution to coding at the average ASA level:** Same analysis as in Fig. 3C-D considering average ASA. Percentage of non-selective neurons at the FSA level that took part in at least one coding assembly, i.e. selective for either of the two variables with its average ASA ($p<0.05$, two-sample t-test). **A-B** Results are presented for both pair assemblies (**A)** and full-size assemblies (**B)** divided according to the number of assemblies to which a neuron belongs. In both cases, there is a significant upward trend in these percentages with the number of assemblies formed by a neuron (Cochran-Armitage test for trend). *** $p<0.001$. For the exact p-values, refer to Table S1.

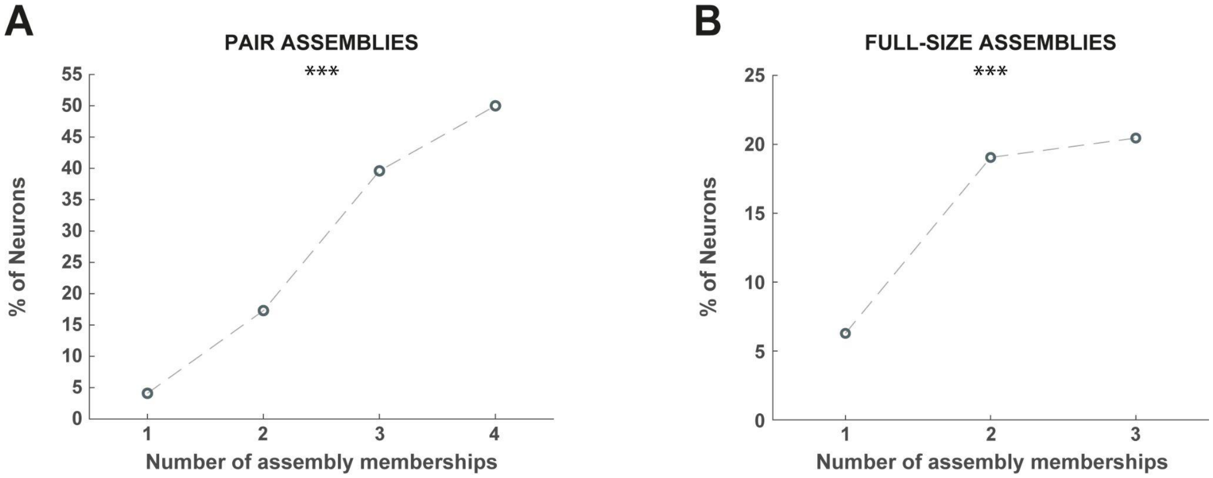


**Figure S3. Multiple coding at the average ASA level:** Same analysis as in Fig. 4C-D considering the average ASA. It shows the percentage of non-multiple selectivity neurons, indicating neurons encoding none or only one of the two variables (response direction or goal color) with their FSA, that took part in at least one assembly coding for each of the two variables considered. **A-B** Results are presented for both pair assemblies (**A)** and full-size assemblies (**B)** divided according to the number of assemblies to which a neuron belongs. These percentages increase significantly with the number of assemblies in which a neuron participates (Cochran-Armitage test for trend) for both pair assemblies and full-size assemblies. *** p<0.001. For the exact p-values, refer to Table S1.

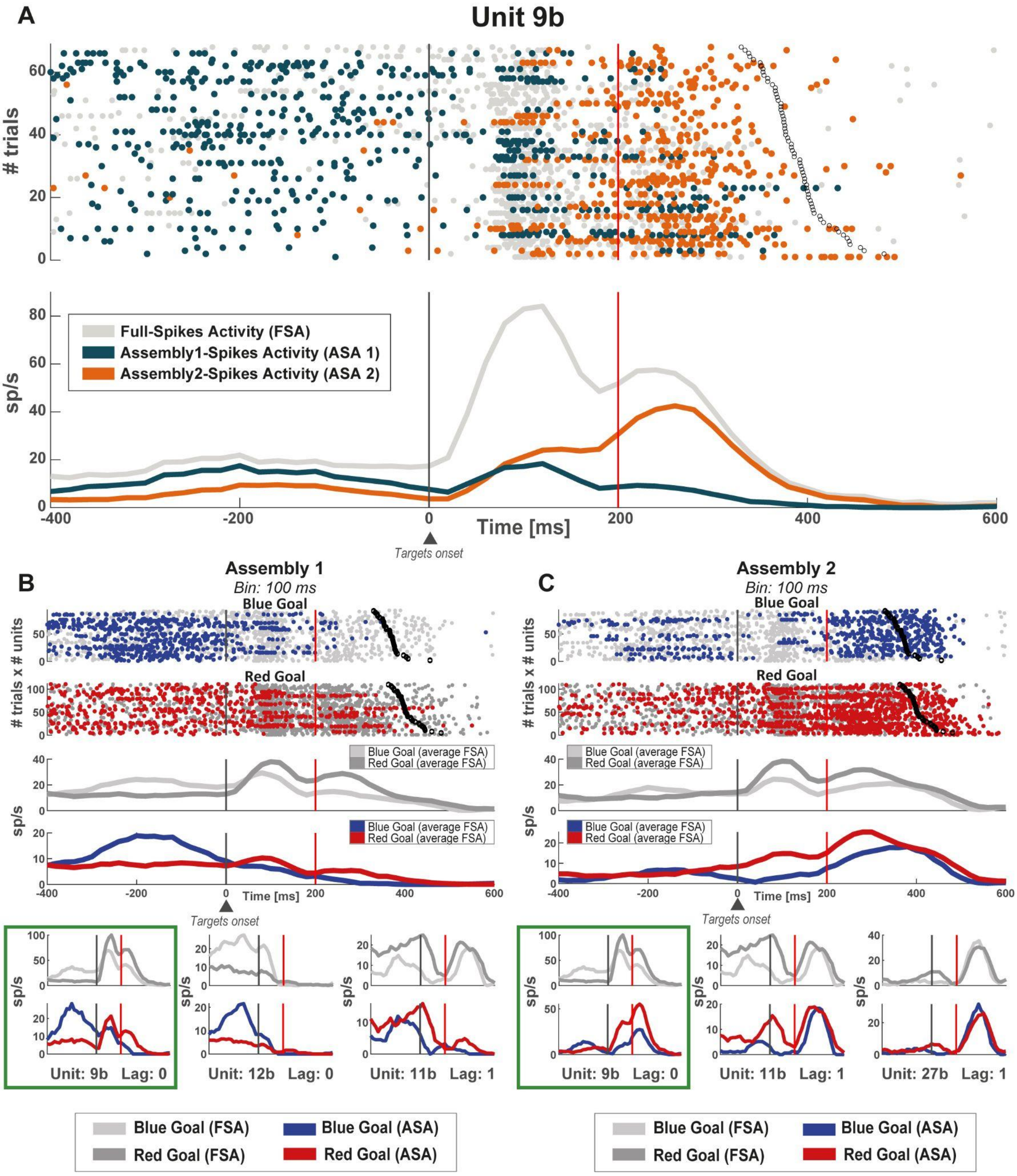


**Figure S4. Activity reconfiguration in a switch neuron. A** Neuron (9b) with an activity reconfiguration between Pre- and Post-go epochs through its participation in different assemblies. **B** At the top, the raster plot and the average FSA and ASA of neurons in Assembly 1 sorted by goal color. At the bottom, the individual mean ASA for the two goal colors of the three neurons composing the Assembly 1 are shown: the first (unit 9b) displayed on the left, as shown in **A**, the second (12b) displayed in the center, and the third (11b) displayed on the right. **C** Same as in (**B)** but for the Assembly 2 with two neurons shared with Assembly 1 (9b, 11b) and a third different neuron (27b) not shared with Assembly 1, displayed on the right. In (**A**-**C**) the black vertical bar indicates the target onset (i.e. the end of the Pre-Go epoch) and the red vertical bar indicates the beginning of the Post-Go epoch, which ends at the response time (black circles). In (**B**) and (**C**) the green boxes indicate the neuron (9b) in (**A**).

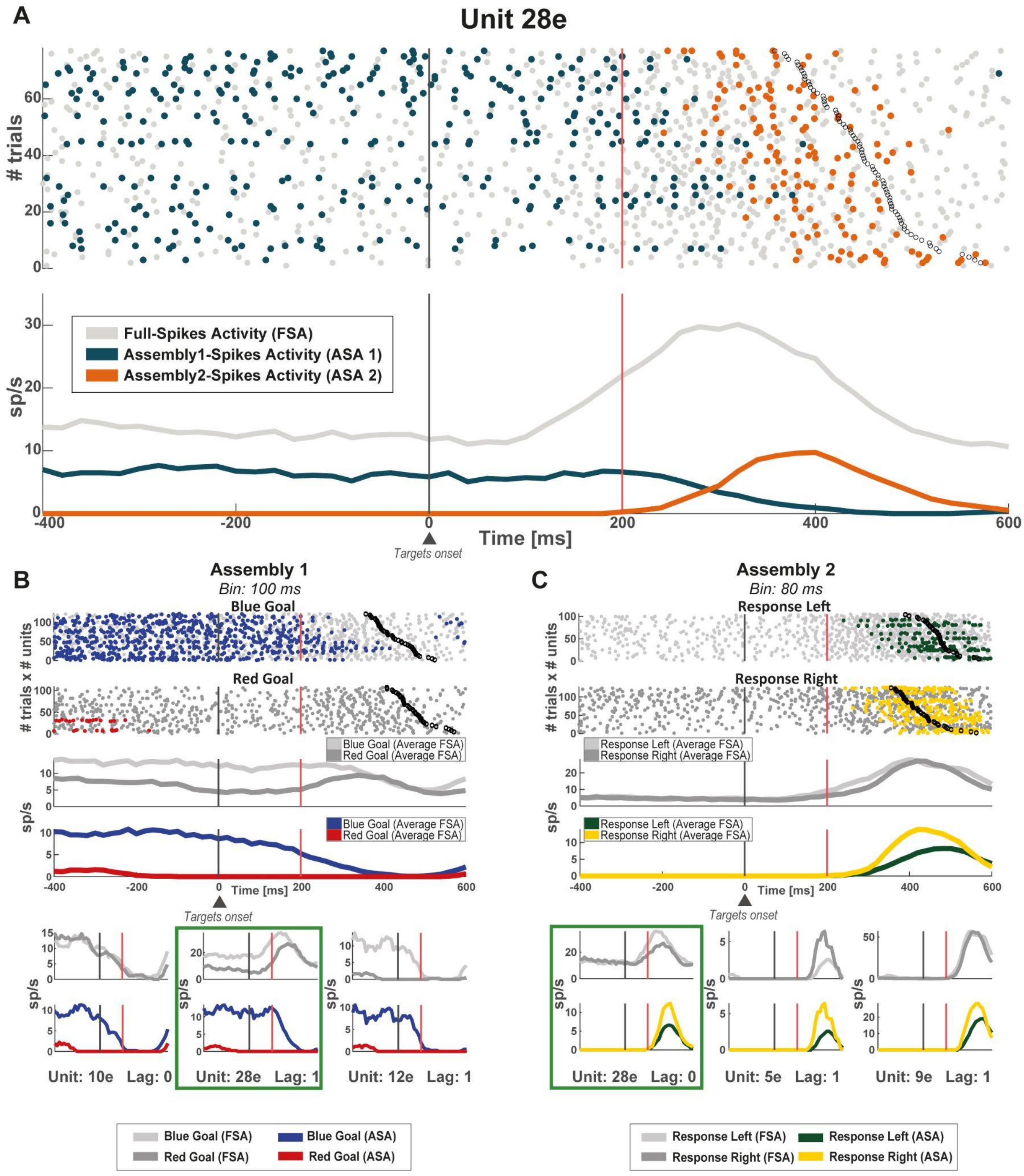


**Figure S5. Activity reconfiguration in a persistent neuron. A** Neuron (28e) with an activity reconfiguration between Pre- and Post-go epochs through its participation in different assemblies. The two assemblies are active in exclusive epochs. **B** At the top, the raster plot is sorted by goal color and the average FSA and ASA of the neurons of the Assembly 1 are shown sorted by each goal. At the bottom, the individual average ASA composing the Assembly 1 for the three neurons, sorted by goal color, that contributed to encode the blue goal are presented: the first on the left (unit 10e), as shown in A, the second (28e), as presented in A, at the center, and the third (12e) on the right. **C** Same as in (**B)** but for the Assembly 2 with one neuron shared with the Assembly 1 (28e), and with two new neurons (5e, 9e) not shared with Assembly 1. All neurons in Assembly 2 shared a preference for the right response. In (A-C) the black vertical bar indicates the target onset (i.e. the end of the Pre-Go epoch), the red vertical bar indicates the beginning of the Post-Go epoch, which ends at the response time (black circles). In (**B**) and (**C**), the green boxes indicate the neuron (28e) in (**A**).

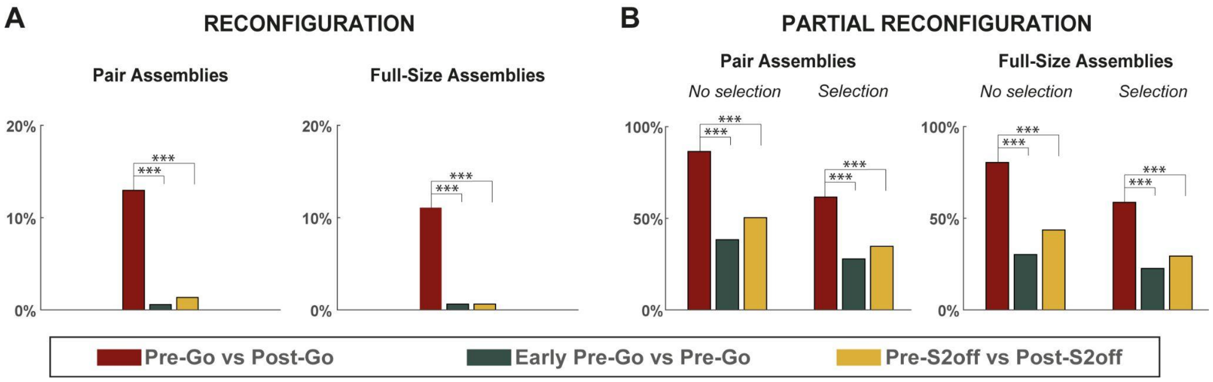


**Fig. S6. Network Reconfiguration at the average ASA level:** Same analysis as in Fig. 6D-E considering the average ASA. **A** Percentage of neurons with activity reconfigurations between each of the three pairs of epochs. The same analysis was performed for both pair assemblies and full-size assemblies. **B** Same as in (**A)** but considering partial activity reconfigurations. We considered two different populations: the entire population of neurons belonging to more than one assembly (*No selection* panel) and the subpopulation with no statistically significant difference in the FSA between each pair of epochs (*Selection* panel). The same analysis was performed for both pair assemblies and full-size assemblies. Statistical significance was assessed using a chi-square test. *** p<0.001. Table S1 reports the exact p-values.

| | SAME PREFERENCE | | | | | | | |
|---|---|---|---|---|---|---|---|---|
| | PAIR ASSEMBLIES | | | | FULL-SIZE ASSEMBLIES | | | |
| | *Left/Right* | | *Blue/Red* | | *Left/Right* | | *Blue/Red* | |
| ***ALL ASSEMBLIES*** | ***FSA*** | ***ASA*** | ***FSA*** | ***ASA*** | ***FSA*** | ***ASA*** | ***FSA*** | ***ASA*** |
| | *3.0e-04* | *2.8e-101* | *0.25* | *2.9e-81* | *0.011* | *1.9e-79* | *0.79* | *2.5e-65* |
| | ***FSA VS ASA*** | | ***FSA VS ASA*** | | ***FSA VS ASA*** | | ***FSA VS ASA*** | |
| | *5.1e-38* | | *6.4e-38* | | *2.2e-31* | | *2.5e-32* | |
| ***ONLY CODING ASSEMBLIES*** | ***FSA*** | ***ASA*** | ***FSA*** | ***ASA*** | ***FSA*** | ***ASA*** | ***FSA*** | ***ASA*** |
| | *5.9e-14* | *3.3e-85* | *3.1e-12* | *4.4e-92* | *2.5e-08* | *1.2e-53* | *5.7e-08* | *8.7e-53* |
| | ***FSA VS ASA*** | | ***FSA VS ASA*** | | ***FSA VS ASA*** | | ***FSA VS ASA*** | |
| | *3.1e-21* | | *1.9e-25* | | *6.4e-16* | | *1.0e-19* | |

| | TRENDS | | | |
|---|---|---|---|---|
| | PAIR ASSEMBLIES | | FULL-SIZE ASSEMBLIES | |
| | ***Becoming Coding*** | ***Becoming Multiple Coding*** | ***Becoming Coding*** | ***Becoming Multiple Coding*** |
| ***ASA*** | *7.2e-05* | *6.7e-24* | *0.0026* | *9.4e-11* |
| ***Average ASA*** | *1.4e-12* | *4.5e-35* | *4.5e-07* | *1.7e-08* |

| | RECONFIGURATION | | | |
|---|---|---|---|---|
| | PAIR ASSEMBLIES | | FULL-SIZE ASSEMBLIES | |
| | ***Pre/Post-Go VS Pre/Post-S2off*** | ***Pre/Post-Go VS EarlyPre/Pre-Go*** | ***Pre/Post-Go VS Pre/Post-S2off*** | ***Pre/Post-Go VS EarlyPre/Pre-Go*** |
| ***ASA*** | *1.6e-07* | *5.0e-08* | *2.8e-04* | *7.5e-05* |
| ***Average ASA*** | *4.5e-13* | *2.3e-15* | *2.3e-08* | *2.3e-08* |

| | PARTIAL RECONFIGURATION | | | |
|---|---|---|---|---|
| | PAIR ASSEMBLIES | | FULL-SIZE ASSEMBLIES | |
| | ***Pre/Post-Go VS Pre/Post-S2off*** | ***Pre/Post-Go VS EarlyPre/Pre-Go*** | ***Pre/Post-Go VS Pre/Post-S2off*** | ***Pre/Post-Go VS EarlyPre/Pre-Go*** |
| ***Selection (ASA)*** | *3.3e-05* | *4.1e-07* | *0.0013* | *1.7e-05* |
| ***Selection*** | *3.9e-06* | *1.2e-09* | *2.6e-05* | *3.3e-08* |

| | | | | |
|---|---|---|---|---|
| ***(Average ASA)*** | | | | |
| ***No Selection (ASA)*** | *1.8e-36* | *6.6e-54* | *8.9e-21* | *2.3e-33* |
| ***No Selection (Average ASA)*** | *1.5e-32* | *1.3e-52* | *4.0e-20* | *1.5e-34* |

**Table S1. P-values from statistical analysis: (Same Preference)** P-values from the statistical analysis of the concordance in preference. The chance level was set to 0.5 when considering pair assemblies, i.e. we considered an equal probability to have the same preference against to have different preferences. In the full-size assemblies, differently from pair assemblies, the probability of having the same preference for all the neurons in each assembly is strictly related to the number of neurons in each assembly. For this reason, the chance level was calculated considering assembly numerosity (see *Methods*). To test the deviation from chance level, we utilized a binomial test. Additionally, we used a chi-square test to compare FSA and ASA. **(Flexibility)** P-values of the Cochran–Armitage test for trend used to assess significance on the increasing trend of the fraction of neurons contributing to coding or to multiple coding only at the assembly level. **(Reconfiguration - Partial Reconfiguration)** P-values of the chi-square test of the comparison of the reconfigurations in the Pre-Go/Post-Go epochs with the two control pairs of epochs.
